# SARS-CoV-2 spike glycoprotein S1 induces neuroinflammation in BV-2 microglia

**DOI:** 10.1101/2020.12.29.424619

**Authors:** Olumayokun A Olajide, Victoria U Iwuanyanwu, Oyinkansola D Adegbola, Alaa A Al-Hindawi

## Abstract

In addition to respiratory complications produced by SARS-CoV-2, accumulating evidence suggests that some neurological symptoms are associated with the disease caused by this coronavirus. In this study, we investigated the effects of the SARS-CoV-2 spike protein S1 stimulation on neuroinflammation in BV-2 microglia. Analyses of culture supernatants revealed an increase in the production of TNFα, IL-6, IL-1β and iNOS/NO. S1 also increased protein levels of phospho-p65 and phospho-IκBα, as well as enhancing DNA binding and transcriptional activity of NF-κB. These effects of the protein were blocked in the presence of BAY11-7082 (1 μM). Exposure of S1 to BV-2 microglia also increased the protein levels of NLRP3 inflammasome and enhanced caspase-1 activity. Increased protein levels of p38 MAPK was observed in BV-2 microglia stimulated with the spike protein S1 (100 ng/mL), an action that was reduced in the presence of SKF 86002 (1 μM). Results of immunofluorescence microscopy showed an increase in TLR4 protein expression in S1-stimulated BV-2 microglia. Furthermore, pharmacological inhibition with TAK 242 (1 μM) and transfection with TLR4 siRNA resulted in significant reduction in TNFα and IL-6 production in S1-stimulated BV-2 microglia. These results have provided the first evidence demonstrating S1-induced neuroinflammation in BV-2 microglia. We propose that induction of neuroinflammation by this protein in the microglia is mediated through activation of NF-κB and p38 MAPK, possibly as a result of TLR4 activation. These results contribute to our understanding of some of the mechanisms involved in CNS pathologies of SARS-CoV-2.

## Introduction

The emergence of the severe acute respiratory syndrome coronavirus-2 (SARS-CoV-2) in 2019 has resulted in a global pandemic affecting most countries and territories. To date, there are 234,809,103 confirmed cases and 4,800,375 deaths [1]. The most common complications of illness due to SARS-CoV-2 infection are mainly related to respiratory symptoms [2–4]. SARS-CoV-2 infection causes significant damage to the alveolus, interstitial and intra-alveolar oedema, infiltration of inflammatory cells (mainly macrophages) [5].

In addition to respiratory complications of SARS-CoV-2 illness, accumulating evidence suggest that neurological symptoms are associated with COVID-19, the disease caused by the novel coronavirus [6]. These symptoms include headaches, loss of smell, confusion and strokes [7]. Furthermore, a retrospective study in China reported neurological symptoms such as cerebrovascular diseases, consciousness impairment, and skeletal muscle symptoms [8]. Interestingly, the SARS-CoV-2 subunit S1 has been shown in a study to promote loss of barrier integrity and trigger a pro-inflammatory response in a 3D model of the human blood-brain barrier [9], suggesting a possible mechanism for the neurological symptoms induced by the coronavirus. Other studies have proposed mechanisms of SARS-CoV-2-mediated CNS dysfunction [10, 11]. Further understanding of the cellular mechanisms involved in neurological consequences of SARS-CoV-2 infection is critical for identifying pharmacological strategies for prevention and treatment.

Microglia are brain-resident macrophages that regulate brain development, maintenance of neuronal networks, and injury repair [12]. In the resting state, microglia survey their surrounding microenvironment through the projection and retraction of their highly motile processes [13]. However, in the presence of injury, or pathology within the CNS they become activated [14]. During neuroinflammation, polarised M1 microglia produce pro-inflammatory cytokines, neurotoxic molecules, such as tumour necrosis factor (TNF)-α, IL-6, IL-1β, nitric oxide (NO), reactive oxygen species (ROS), which contribute to neuronal dysfunction, while polarised M2 microglia secrete anti-inflammatory mediators such as IL-10 and transforming growth factor (TGF-β), which are involved in restoring homeostasis [15–18]. Microglia-mediated neuroinflammation therefore plays significant roles in many neurological [19–23] and neuropsychiatric disorders [24–27]. Based on evidence linking SARS-CoV-2 infection with neurological symptoms, we carried out investigations to determine the effects of a recombinant SARS-CoV-2 spike protein S1 sub-unit on the release of pro-inflammatory mediators in BV-2 microglia.

## Materials and methods

### Materials

Recombinant human coronavirus SARS-CoV-2 spike protein S1 (Accession <u>MN908947</u>) was purchased from Abcam. The protein was suspended in sterile water for functional studies. The following drugs were used: BAY11-7082 (Sigma), CRID3 sodium salt (Tocris), SKF 86002 dihydrochloride (Tocris) and TAK 242 (Tocris).

### Cell culture

BV-2 mouse microglia cell line (ICLCATL03001) was purchased from Interlab Cell Line Collection (Banca Biologica e Cell Factory, Italy) and cultured in RPMI medium supplemented with 10% foetal bovine serum.

### Determination of cytokine production

Initial pilot experiments were conducted to evaluate effects of 10, 50, 100, 500 and 1000 ng/mL of S1 on TNFα production in BV-2 microglia seeded out in 24-well plates at 5 × 10^4^ cells/mL. These concentrations of the protein were used to stimulate the cells for 1, 3, 6, 12 and 24 h. Levels of TNFα in culture supernatants were then determined using mouse instant ELISA™ kit (Thermo Scientific).

Subsequent experiments employed (10, 50 and 100 ng/mL) of S1 for determining effects on TNFα, IL-6 and IL-1β. In this case, cultured BV-2 microglia were seeded out in 24-well plate at 5 × 10^4^ cells/mL and treated with protein S1 (10, 50 and 100 ng/mL) for 24 h. Thereafter, medium was collected and centrifuged to obtain culture supernatants. Levels of TNFα were determined as described above, while levels of IL-6 in supernatants were determined using IL-6 mouse ELISA kit (Thermo Scientific). Similarly, levels of IL-1β were evaluated using IL-1β mouse ELISA kit (Thermo Scientific).

### Measurement of nitric oxide (NO) production

The Griess reagent (Promega) was used to determine the levels of levels of nitric oxide (NO) in supernatants of cultured BV-2 microglia treated with protein S1 (10, 50 and 100 ng/mL) for 24 h. Following treatment with S1, Griess assay (which measures nitrite, a stable breakdown product of NO) was carried out by adding 50 μL of sulphanilamide to 50 μL of culture supernatants, followed by incubation in the dark at room temperature for 10 min. Thereafter 50 μL of N-1-naphthyl ethylenediamine dihydrochloride (NED) was added and the mixture incubated for a further 10 min. Absorbance was measured at 540nm using a Tecan Infinite M microplate reader. Nitrite concentrations were determined from a nitrite standard reference curve.

### In cell-western/cytoblot

The in-cell western is a proven method for the rapid quantification of proteins in cells [28, 29]. BV-2 microglia were seeded into a 96-well plate at 5 × 10^4^ cells/mL, and cells treated at 70% confluence. At the end of each experiment, cells were fixed with 8% formaldehyde solution (100 μL) for 15 min., followed by washing with PBS. The cells were then incubated with primary antibodies overnight at 4°C. The following antibodies were used: rabbit anti-iNOS (Abcam), rabbit anti-phospho-p65 (Cell Signalling Technology), rabbit anti-phospho-IκBα (Santa Cruz Biotechnology), rabbit anti-NLRP3 (Abcam), and rabbit anti-phospho-p38 (Cell Signalling Technology) antibodies. Thereafter, cells were washed with PBS and incubated with anti-rabbit HRP secondary antibody for 2 h at room temperature. Then, 100 μL HRP substrate was added to each well and absorbance measured at 450nm with a Tecan Infinite M microplate reader. Readings were normalised with Janus Green normalisation stain (Abcam).

### NF-κB luciferase reporter gene assay

BV-2 cells were seeded in 24-well plate at a density of 4 × 10^4^ cells/mL. At 60% confluence, RPMI medium was replaced with Opti-MEM, with a further incubation for 2 h at 37°C. Transfection of BV-2 cells was performed by preparing a Glia-Mag transfection reagent (OZ Biosciences) and Cignal NF-κB luciferase reporter (Qiagen) complex at a ratio 3:1 in 50 μL Opti-MEM. The complex was added to BV-2 cells, and the plate placed on a magnetic plate (OZ Biosciences) and incubated at 37°C for 30 min, followed by a further magnetic plate-free incubation for 2 h.

Thereafter, medium was changed to serum-free RPMI and cells treated with spike S1 (10, 50 and 100 ng/mL) for 6 h. This was followed by a Dual-Glo^®^;reporter assay (Promega). Firefly and renilla luminescence were measured using a FLUOstar OPTIMA microplate reader (BMG Labtech).

### NF-κB transcription factor binding assay

The NF-κB p65 transcription factor assay is a non-radioactive ELISA-based assay for evaluating DNA binding activity of NF-κB in nuclear extracts. BV-2 microglia were seeded in a 6-well plate at a density of 4 × 10^4^ cells/mL. The cells were then incubated with 10, 50 and 100 ng/mL of S1 for 60 min. At the end of the incubation, nuclear extracts were prepared from the cells and subjected to NF-κB transcription factor binding assay according to the instructions of the manufacturer (Abcam).

### Caspase-Glo^®^1 inflammasome assay

The caspase-Glo^®^1 inflammasome assay (Promega) was used to measure the activity of caspase-1 directly in live cells or culture supernatants [30]. BV-2 microglia were seeded out in 24-well plate at a density of 4 × 10^4^ cells/mL and stimulated with S1 (10, 50 and 10 ng/mL) for 24 h. After incubation, cell culture supernatants were collected and mixed with equal volume of Caspase-Glo^®^ 1 reagent or Caspase-Glo^®^ 1 reagent + YVAD-CHO (1 μM) in a 96-well plate. The contents of the wells were mixed using a plate shaker at 400rpm for 30 seconds. The plate was then incubated at room temperature for 60 min, followed by luminescent measurement of caspase-1 activity with a FLUOstar OPTIMA reader (BMG LABTECH).

### Effects of BAY11-7082, CRID3 and SKF 86002 on S1-induced neuroinflammation

BAY11-7082 is an anti-inflammatory small molecule inhibitor of NF-κB, which has also been proposed as a potential disease-modifying strategy to combat the severity of COVID-19 [31]. CRID3 is an NLRP3 inflammasome inhibitor, which has been shown to reduce interleukin-1β (IL-1β) production *in vivo* and attenuated the severity of experimental autoimmune encephalomyelitis [32]. SKF 86002 is a bicylic pyridinylimidazole which has been widely reported to inhibit cytokine production by targeting p38 MAPK [33].

Consequently, the effects of these inhibitors on the release of TNFα, IL-6 and IL-1β production were investigated by pre-treating BV-2 microglia with BAY11-7082 (1 μM), CRID3 (1 μM), and SKF 86002 (1 μM) 60 min prior to stimulation with S1 (100 ng/mL) for a further 24 h. Culture supernatants were collected and analysed for levels of TNFα, IL-6 and IL-1β using mouse ELISA kits (Thermo Scientific).

### Immunofluorescence microscopy

Immunofluorescence microscopy was carried out by seeding out BV2 microglia in a 24-well cell imaging plate (Eppendorf) at a density of 5 × 10^4^ cells/ml. Thereafter, cells were stimulated with S1 (100 ng/ml) for 24 h. After stimulation, cells were washed with PBS and fixed with 500 μL formaldehyde (4%) for 15 min at room temperature. This was followed by PBS wash and blocking with 500 μL of 5% Bovine Serum Albumin for 60 min at room temperature. Cells were then incubated overnight at 4°C with either Iba-1 Alexa Fluor^®^ 488 antibody (Santa Cruz Biotechnology; 1:500) or TLR4 antibody (Santa Cruz Biotechnology; 1:500). In experiments employing unconjugated primary antibody, cells were washed with PBS and further incubated in the dark for 2 h with Alexa Fluor^®^ 488-conjugated donkey anti-rabbit IgG secondary antibody (Thermo Scientific; 1:500). Then, cells were stained with 50 nM of 4′, 6 diamidino-2-phenylindole dihydrochloride (DAPI; Invitrogen) for 3 min and washed. Images were captured using EVOS FLoid^®^ cell imaging station (Invitrogen).

### Effects of TAK 242 on spike protein S1-induced increased production of pro-inflammatory mediators

TAK 242 is a small molecule inhibitor of inflammation which has been reported to reduce nitric oxide and cytokine production in LPS stimulated mouse macrophages, and in a mouse model [34]. The anti-inflammatory activity of TAK 242 has been confirmed to be mediated through inhibition of TLR4 signalling [35]. The effects of TAK 242 were therefore investigated on S1-induced increased production of TNFα, IL-6, IL-1β. BV-2 microglia were pre-treated with TAK 242 (1 μM). After 60 min, cells were stimulated with S1 (100 ng/mL) for a further 24 h. Levels of TNFα and IL-6 in culture supernatants were measured using mouse ELISA kits (Thermo Scientific).

### siRNA-mediated knockdown of TLR4

In order to further confirm the possible roles of TLR4 in S1-induced increased production of pro-inflammatory mediators, experiments were carried out to knockdown TLR4 in BV-2 microglia. BV-2 cells were seeded out into 6-well plates at concentration 4 × 10^4^ cells/ml and cultured until 60% confluence. Thereafter, culture medium in each well was changed to Opti-MEM (1000 μL) followed by incubation for 2 h at 37°C. Complexes containing 1.8 μL of Glial-Mag transfection reagent (OZ Biosciences) and 2 μL of TLR4 or control siRNA (Santa Cruz Biotechnology) were then added to the cells. The culture plate was placed on a magnetic plate (OZ Biosciences) for 30 min, followed by incubation at 37°C for a further 24 h. Following successful transfection, the effects of loss-of-function of TLR4 were evaluated by stimulating the cells with SARS-CoV-2 spike protein S1 (100 ng/mL) for 24 h. Culture supernatants were collected and analysed for levels of TNFα and IL-6 using mouse ELISA kits**Statistical analyses**

Data are expressed as mean ± SEM for at least three independent experiments (n=3) and analysed using one-way analysis of variance (ANOVA) with post hoc Tukey’s test (for multiple comparisons). Statistical analysis were conducted using the GraphPad Prism software (version 9).

## Results

### Effects of S1 on pro-inflammatory cytokine release in BV-2 microglia

Results of pilot experiments to determine effects of stimulation with 10, 50, 100, 500 and 1000 ng/mL of S1 on the production of TNFα in BV-2 microglia at 1, 3, 6, 12 and 24 h are shown in *Supplementary Data*. Stimulation with all concentrations of the spike protein resulted in a steady increase in the production of TNFα from 1 h post-stimulation, with the maximum release observed 24 h post-stimulation.

Furthermore, there was a concentration-dependent increase in S1-induced increase in TNFα production, which reached a maximum at 100 ng/mL at 3, 6, 12 and 24 h post-stimulation. When compared with effects of 100 ng/mL of S1, there were no further significant increases in TNFα production when the concentration of S1 was increased to 500 and 1000 ng/mL Consequently, in subsequent confirmatory experiments 10, 50 and 100 ng/mL of S1 were then incubated with BV-2 cells for 24 h. Results show that with 10 ng/mL of S1, there was an insignificant (p<0.05) elevation in the secretion of TNFα. However, on increasing the concentration to 50 ng/mL, significant (p<0.05) increase in the production of TNFα was observed (Figure 1A). On increasing the concentration of the protein to 100 ng/mL, there was a ∼15-fold increase in the levels of TNFα in culture supernatants (Figure 1A).

**Figure 1.**
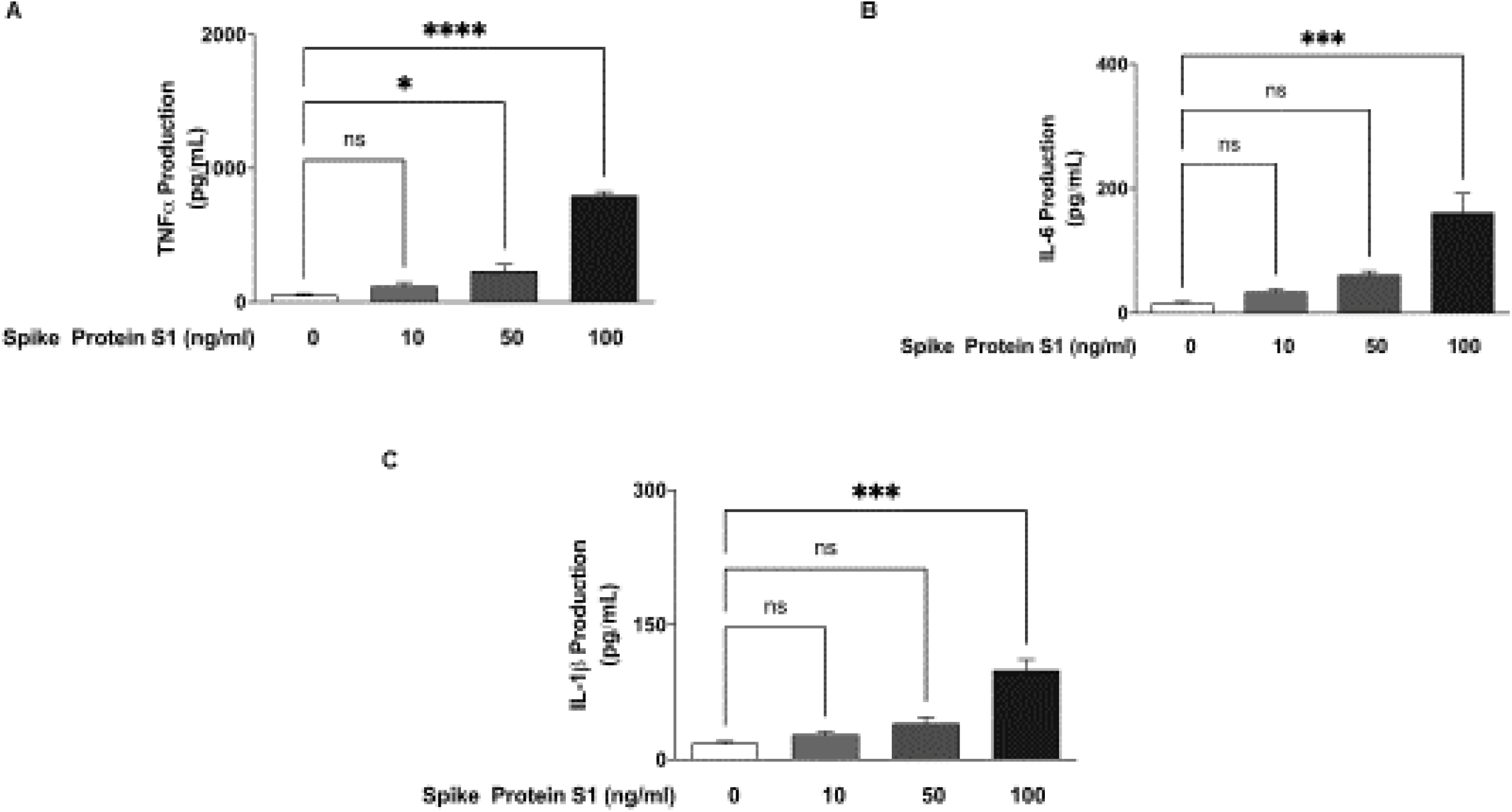
S1 increased levels of TNFα (A), IL-6 (B) and IL-1β (C) in BV-2 microglia after a 24-h incubation. All values are expressed as mean ± SEM for three independent experiments. Data were analysed using One-way ANOVA followed by a post hoc Tukey’s multiple comparison test. *p < 0.05, ***p < 0.001, ****p<0.0001, ns (not significant at p<0.05), compared with untreated control.

### S1 induced an increase in IL-6 and IL-1β production

Results in Figure 1B show that release of IL-6 was not significantly elevated in BV-2 microglia treated with 10 and 50 ng/mL of S1, in comparison with untreated (control) cells. However, incubation of the cells with 100 ng/mL of S1 resulted in ∼10.2-fold and significant (p<0.001) increase in the secretion of IL-6 into culture medium. Similar observations were made in experiments to determine the effects of S1 sub-unit on IL-1β production; at 10 and 50 ng/mL of S1, there was no significant effect on the release of IL-1β. A ∼5.5-fold elevation of IL-1β levels was however observed with 100 ng/mL of S1 (Figure 1C).

### S1 induced an increase in NO production and iNOS protein expression in BV-2 microglia

Analyses of culture supernatants obtained from BV-2 microglia incubated with S1 (10 ng/mL) showed insignificant (p<0.05) increase in the levels of NO, when compared with untreated (control) cells. We further showed that incubation with 50 and 100 ng/mL of S1 caused ∼5.1 and 8.7 increase in NO production, respectively (p<0.01) (Figure 2A).

**Figure 2.**
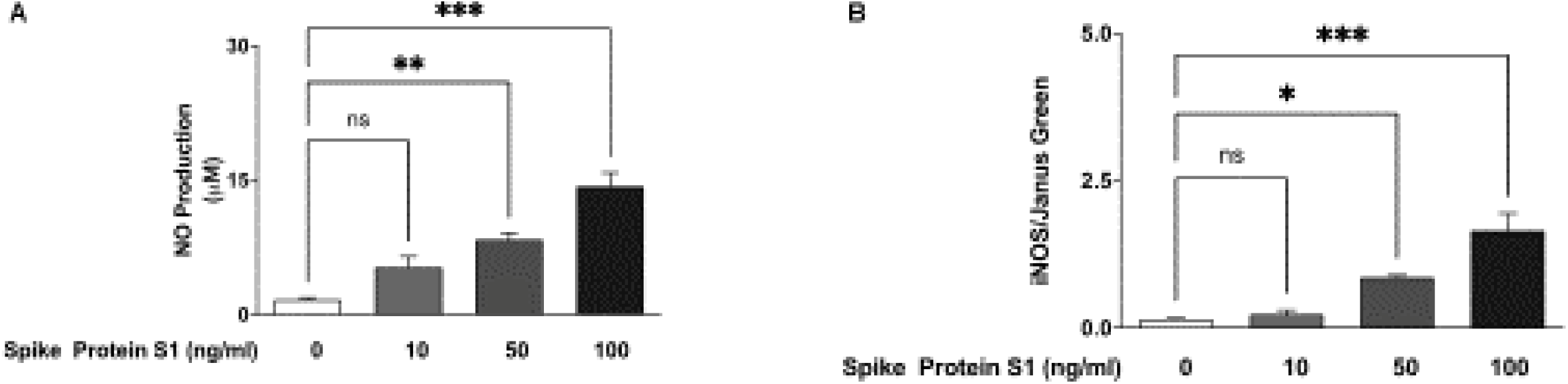
S1 increased levels of NO (A) and iNOS protein (B) in BV-2 microglia after a 24-h incubation. NO levels in culture supernatants were measured using the Griess assay, while in-cell western (cytoblot) assay was used to detect iNOS protein expression. All values are expressed as mean ± SEM for three independent experiments. Data were analysed using One-way ANOVA followed by a post hoc Tukey’s multiple comparison test. *p<0.05, **p < 0.01, ***p < 0.001, ns (not significant at p<0.05), compared with untreated control.

We also used in cell western analyses to further demonstrate that incubation with S1 (50 and 100 ng/mL) resulted in significant (p<0.05) increase in protein levels of iNOS, in comparison with unstimulated cells (Figure 2B).

### S1 increased Iba-1 expression in BV-2 microglia

Immunofluorescence microscopy revealed that unstimulated BV-2 microglia express low levels of Iba-1 protein. However, on activation with S1 (10-100 ng/mL), there were marked increases in expression of (Figure 3).

**Figure 3.**
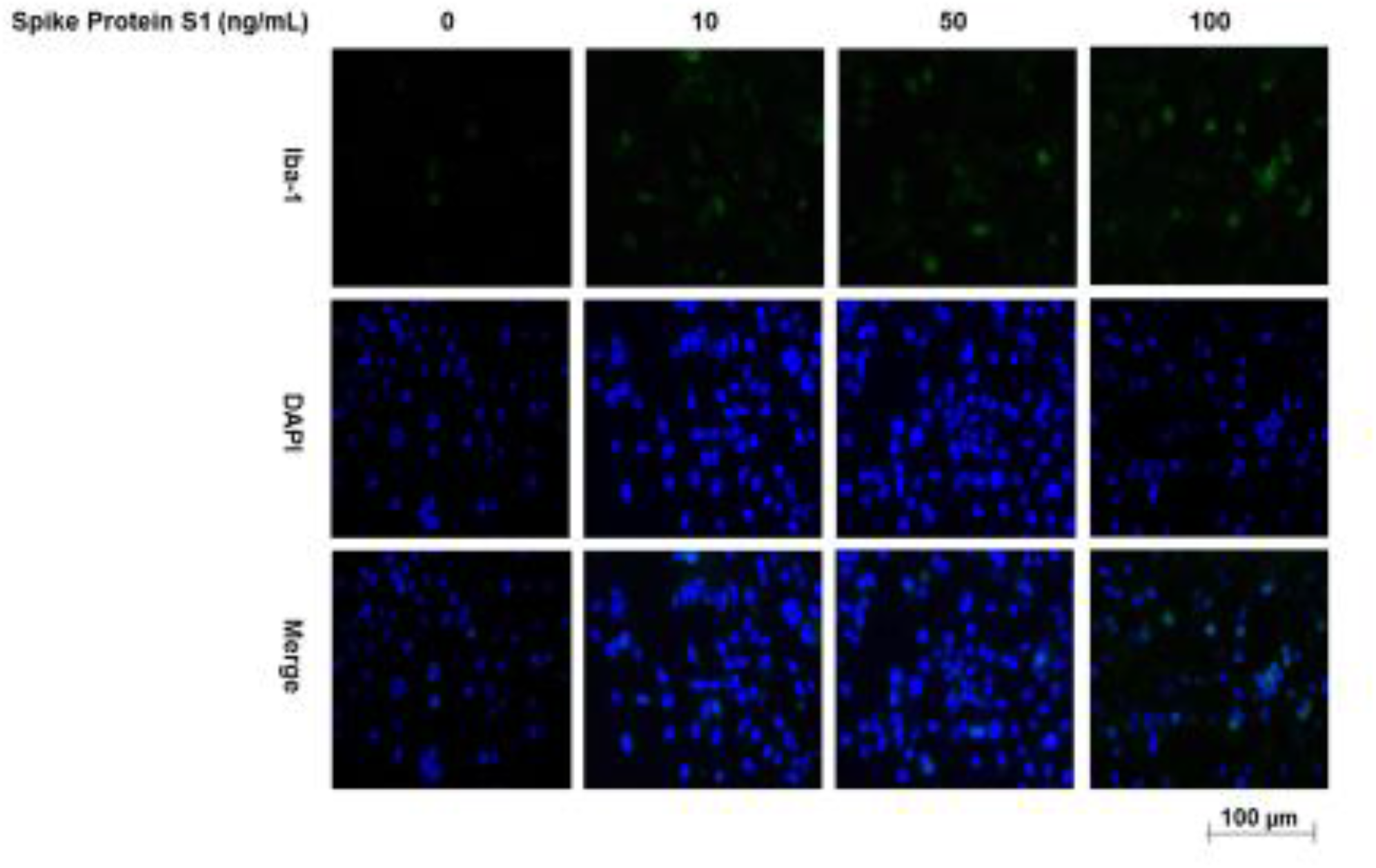
Immunofluorescence microscopy showing increase in Iba-1 protein expression in BV-2 microglia incubated with 10, 50 and 100 ng/mL of S1.

### Spike protein S1 activated NF-κB in BV-2 microglia

Following our observation that S1 induced a significant increase in the production of pro-inflammatory mediators at 100 ng/mL, we were next interested in evaluating the role of NF-κB in its activity at this concentration. Firstly, we used in cell western assays to investigate effects of the protein on the cytoplasmic activation of p65 and IκBα. Results show that S1 (100 ng/mL) significantly (p<0.01) increased protein levels of both phospho-p65 and phospho-IκBα following incubation with BV-2 microglia for 15 min (Figures 4A and 4B). These results also show that protein levels of phospho-p65 and phospho-IκBα were reduced in the presence of the NF-κB inhibitor, BAY11-7082 (1 μM).

**Figure 4.**
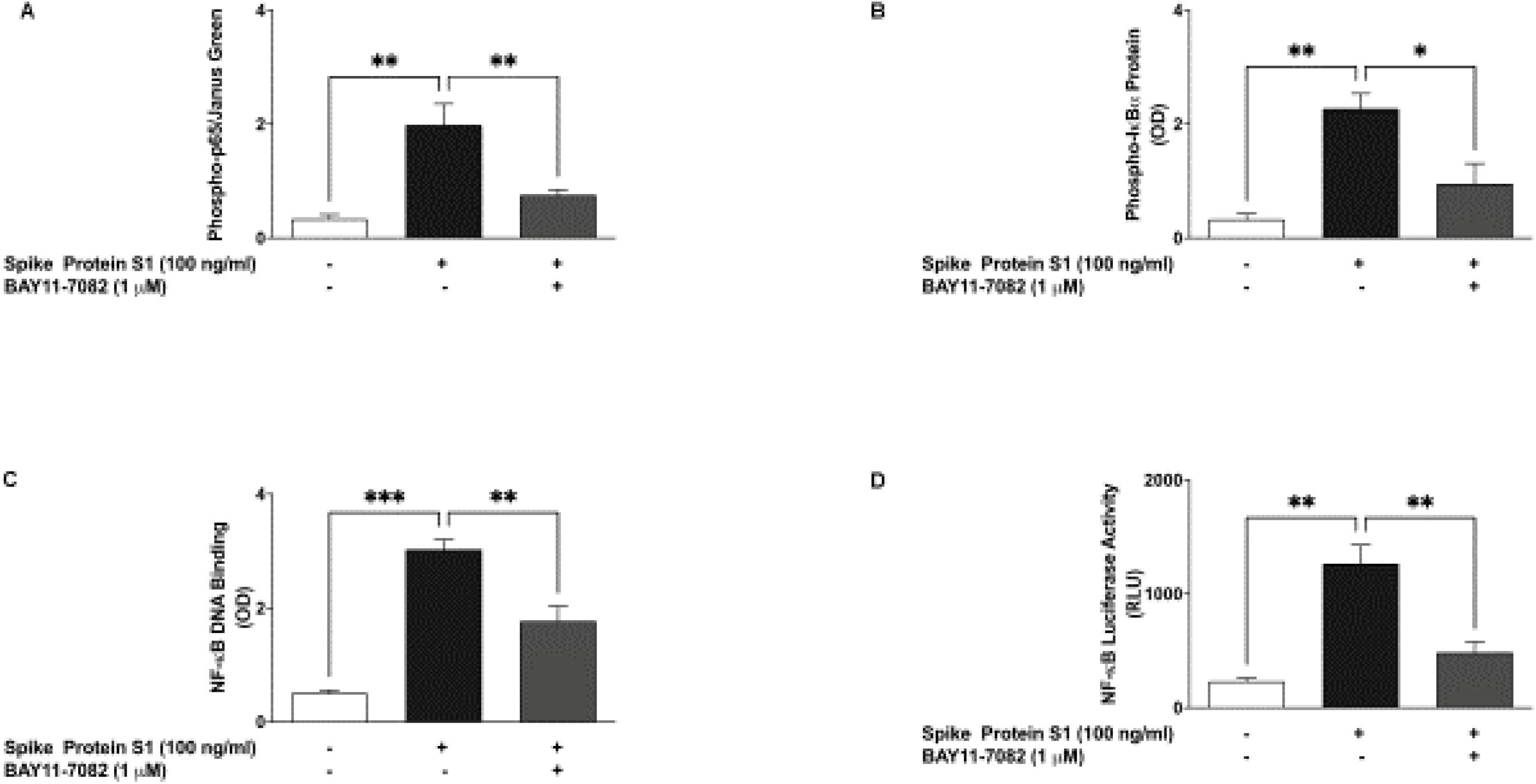
S1 activated NF-κB signalling in BV-2 microglia. Nuclear extracts were collected from cells stimulated with S1 (100 ng/mL) in the absence or presence of BAY11-7082 (1 μM) for 60 min and subjected to DNA binding assays (A). Results of luciferase reporter gene assay showing increased NF-κB transcriptional activity by S1 and its inhibition by BAY11-7082 (1 μM) (B). In-cell western (cytoblot) analyses showing increased protein expressions of phospho-p65 sub-unit (C) and phospho-IκBα (D) following stimulation with S1 (100 ng/mL) for 15 min and inhibition by BAY11-7082 (1 μM). Data were analysed using One-way ANOVA followed by a post hoc Tukey’s multiple comparison test. *p<0.05, **p < 0.01, ***p < 0.001, ns (not significant at p<0.05), compared with untreated control, or S1 stimulation alone.

Results of an NF-κB p65 transcription factor assay revealed that following treatment of BV-2 microglia with S1 (100 ng/mL) for 60 min, there was a significant (p<0.001) increase in the binding of NF-κB in nuclear extracts to consensus-binding sites in the DNA. Interestingly, pre-treatment of BV-2 cells with BAY11-7082 (1 μM) prior to stimulation with S1 produced an inhibition in binding to a double stranded DNA sequence containing the NF-κB response element (Figure 4C). Luciferase reporter gene assays to evaluate the effects of S1 on transcriptional activity showed that S1 stimulation of BV-2 microglia that were transfected with NF-κB plasmid vector resulted in a significant (p<0.01) increase in luciferase activity, in comparison with untreated transfected cells. We further showed that on pre-treating the cells with BAY11-7082 (1 μM), S1-induced increase in NF-κB luciferase activity was markedly reduced (Figure 4D).

### Pre-treatment with BAY11-7082 reduced S1-induced production of pro-inflammatory mediators

In order to further confirm the roles of NF-κB in S1-induced neuroinflammation, BV-2 microglia were pre-treated with BAY11-7082 prior to stimulation with the protein for 24 h, followed by measurements of pro-inflammatory cytokines (TNFα, IL-6 and IL-1α) release in culture supernatants. In the presence of BAY11-7082 (1 μM), S1-induced increases in TNFα, IL-6 and IL-1α production in BV-2 microglia were reduced by ∼59.6%, ∼40.6% and ∼32.8%, respectively (Figures 5A, 5B and 5C).

**Figure 5.**
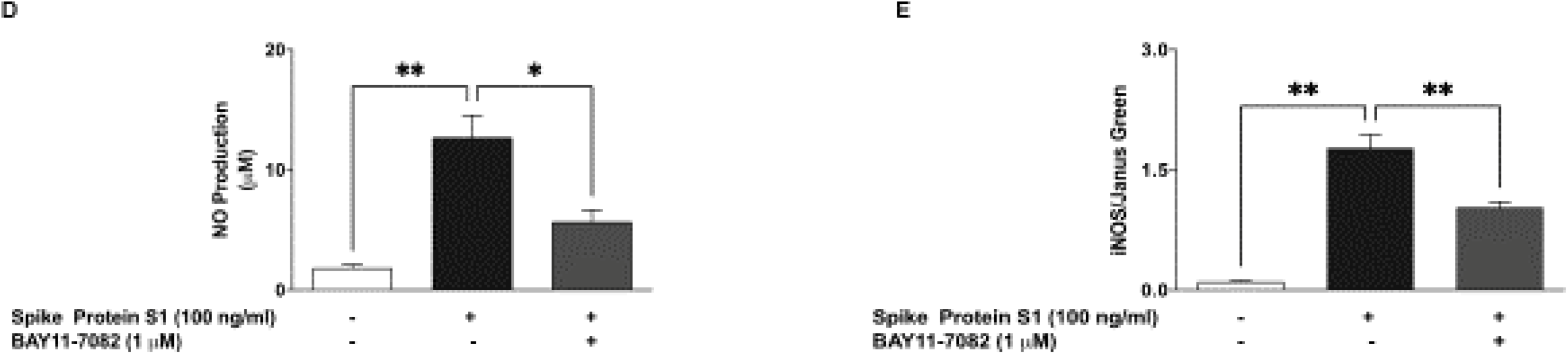
Pre-treatment with BAY11-7082 (1 μM) resulted in inhibition of S1-induced increased production of TNFα (A), IL-6 (B) and IL-1β (C) in BV-2 microglia. Culture supernatants were analysed using ELISA following stimulation for 24 h. Data were analysed using One-way ANOVA followed by a post hoc Tukey’s multiple comparison test. **p < 0.01, ****p<0.0001, compared with untreated control, or S1 stimulation alone. Pre-treatment with BAY11-7082 (1 μM) resulted in inhibition of S1-induced increased production of NO (D), iNOS protein (E) in BV-2 microglia, following stimulation for 24 h. Culture supernatants were analysed using Griess assay, in-cell western (cytoblot) was used for iNOS detection. Data were analysed using One-way ANOVA followed by a post hoc Tukey’s multiple comparison test. *p<0.05, **p < 0.01, ***p<0.001, ****p<0.0001, compared with untreated control, or S1 stimulation alone.

Results in Figures 5D and 5E also show that S1-induced increase in NO production and iNOS protein expression were significantly (p<0.01) reduced when the cells were treated with BAY11-7082 (1 μM) prior to S1 stimulation.

### S1 triggers activation of NLRP3 inflammasome/caspase-1 in BV-2 microglia

The NLRP3 inflammasome is known to contribute to the secretion of IL-1β during inflammation. Encouraged by results of experiments showing an increase in the production of this cytokine following BV-2 stimulation with S1, we next investigated its effect on protein levels of NLRP3 in the presence and absence of known NLRP3 inhibitors, CRID3 and BAY11-7082. Cytoblot analyses revealed that stimulation of BV-2 microglia with S1 (100 ng/mL) for 6 h resulted in ∼11-fold increase in levels of NLRP3 protein. We also observed that S1-induced increase in NLRP3 protein levels were significantly reduced (p<0.05) in the presence of CRID3 (1 μM) and BAY11-7082 (1 μM) (Figure 6A).

**Figure 6.**
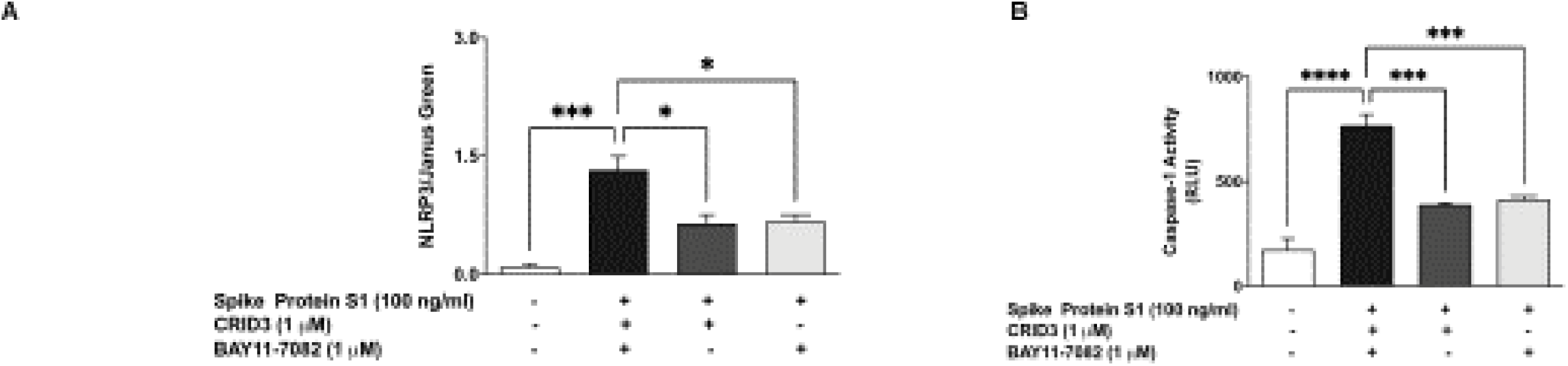
(A) Stimulation of BV-2 microglia with S1 (100 ng/mL) increased protein expression of NLRP3 inflammasome, and was inhibited in the presence of CRID3 (1 μM) and BAY11-7082 (1 μM). (B) Increased caspase-1 activity by S1 (100 ng/mL) BV-2 microglia was reduced in the presence of CRID3 (1 μM) and BAY11-7082 (1 μM). Data were analysed using One-way ANOVA followed by a post hoc Tukey’s multiple comparison test. *p<0.05, **p < 0.01, ***p<0.001, ****p<0.0001, compared with untreated control, or S1 stimulation alone.

We next determined measured caspase-1 activity in culture media of BV-2 cells that were stimulated with S1 for 6 h. Results in Figure 6B show a ∼4.4-fold increase in caspase-1 activity as a result of S1 stimulation. Furthermore, both CRID3 (1 μM) and BAY11-7082 (1 μM) produced significant (p<0.001) inhibition in S1-induced increased caspase-1 activity.

### CRID3 prevented S1-induced increase in IL-1β production

Experiments to evaluate the effects of CRID3 on S1-induced increased secretion of TNFα, IL-6 and IL-1β in BV-2 microglia revealed that pre-treatment with CRID3 (1 μM) was ineffective in reducing TNFα (Figure 7A) and IL-6 (Figure 7B) production. However, pre-treatment of S1-stimulated BV-2 cells with CRID3 significantly (p<0.001) reduced IL-1β production (Figure 7C). We also observed that S1-induced NO production was not reduced by pre-treatment with CRID3 (Figure 7D).

**Figure 7.**
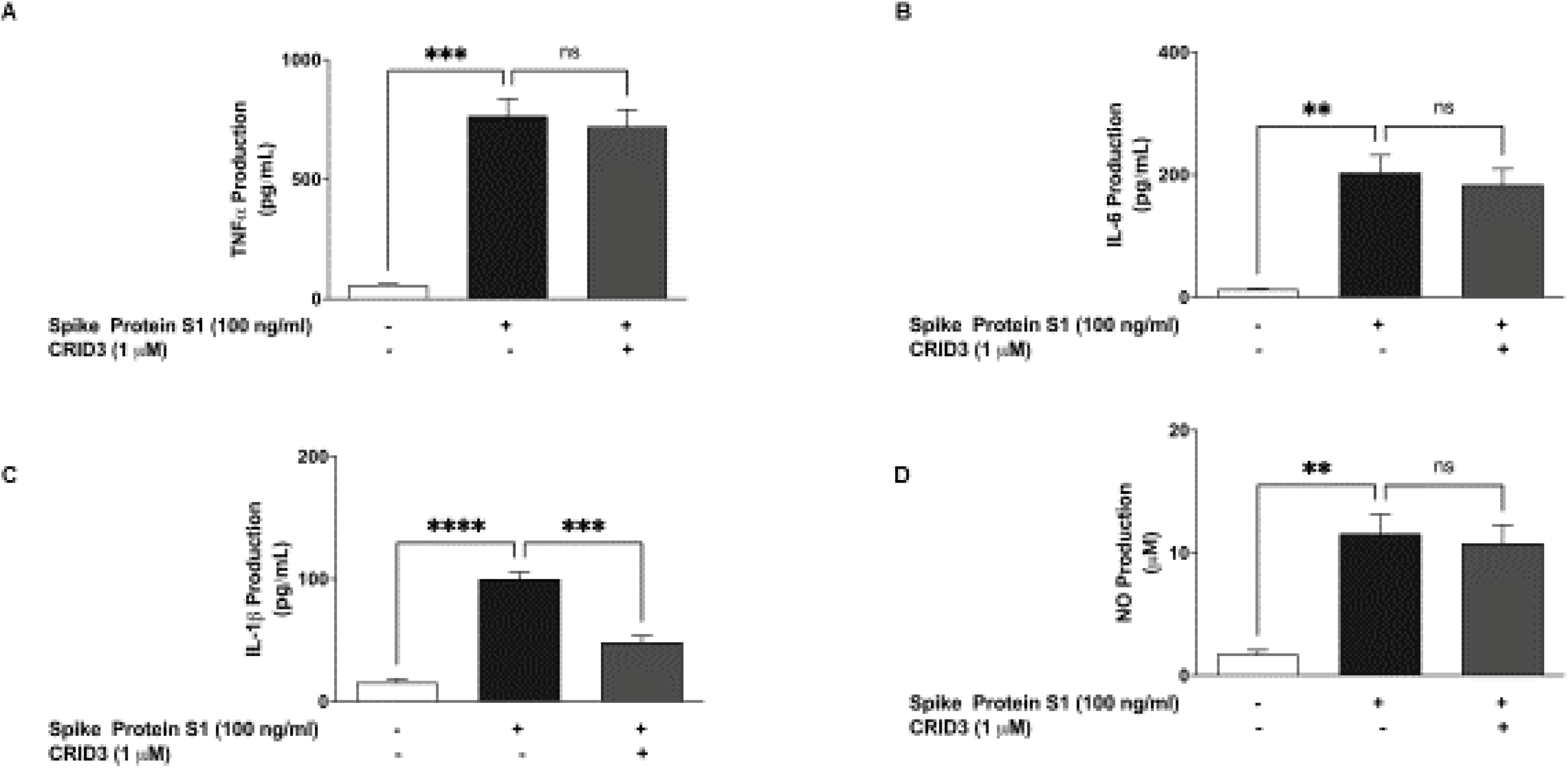
Pre-treatment with CRID3 (1 μM) did not prevent S1-induced increased production of TNFα (A), IL-6 (B), NO (D) while IL-1β production was reduced (C) in BV-2 microglia. Culture supernatants were analysed using ELISA following stimulation for 24 h. Data were analysed using One-way ANOVA followed by a post hoc Tukey’s multiple comparison test. **p < 0.01, ***p<0.001, ****p<0.0001, ns (not significant at p<0.05), compared with untreated control, or S1 stimulation alone.

### S1 activated p38 MAP kinase in BV-2 microglia

Incubation of spike protein S1 (100 ng/mL) with BV-2 microglia for 60 min resulted in a significant (p<0.01) and ∼11.3-fold increase in protein levels of phospho-p38; an outcome that was prevented by pre-treating the cells with SKF86002 (1 μM) for 60 min prior to S1 stimulation (Figure 8A).

**Figure 8.**
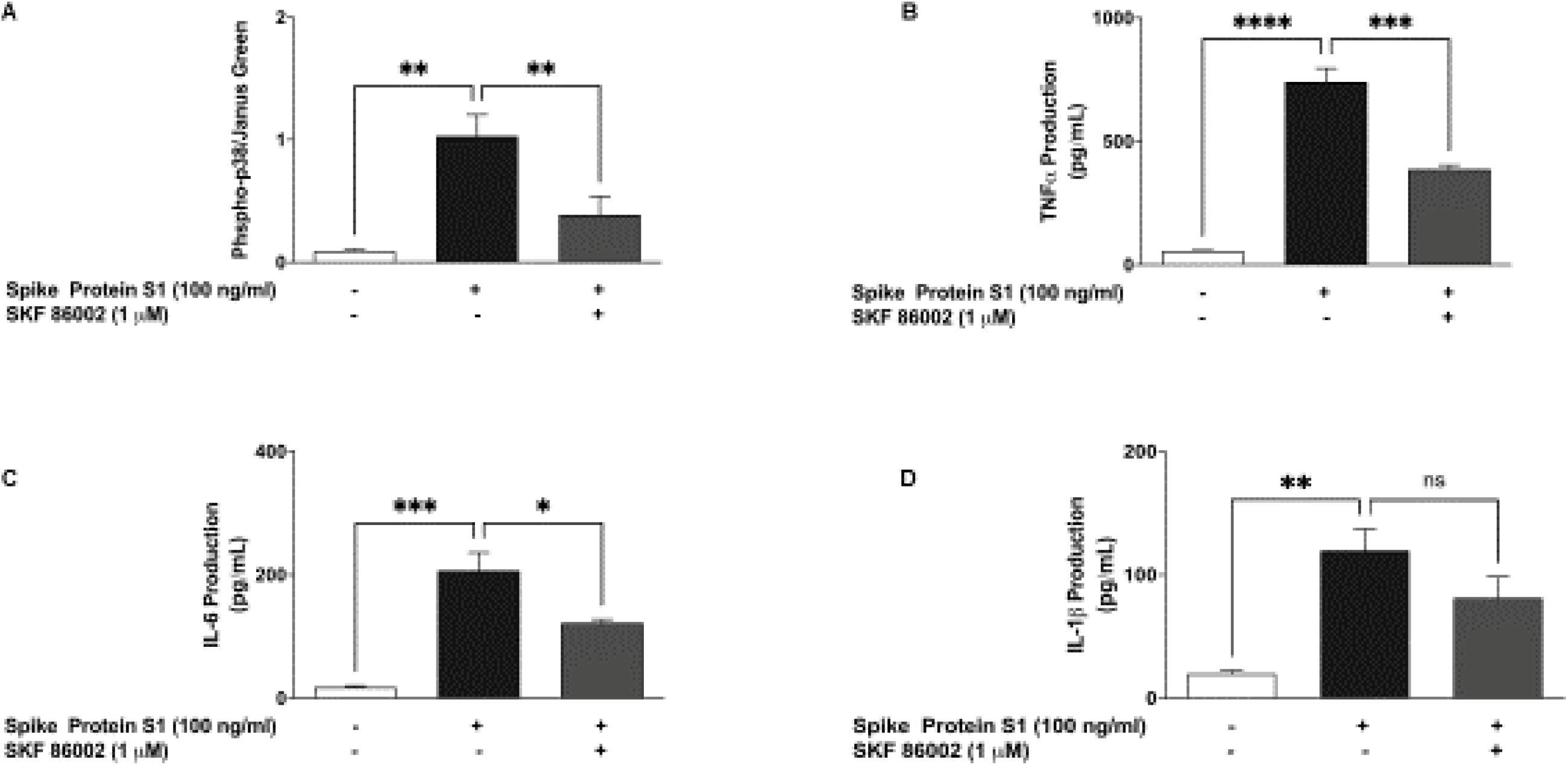
Stimulation of BV-2 microglia with S1 (100 ng/mL) increased protein expression of phospho-p38 MAPK, which was inhibited in the presence of SKF 86002 (1 μM). Data were analysed using One-way ANOVA followed by a post hoc Tukey’s multiple comparison test. *p<0.05, **p < 0.01, compared with untreated control, or S1 stimulation alone.

Based on the results showing that S1 caused an increase in protein levels of phospho-p38 and its inhibition by SKF86002, we were next interested to determine the effects of this inhibitor on cytokine production in S1-stimulated BV-2 microglia. In the presence of SKF86002 (1 μM), TNFα production was reduced by ∼47.9% compared with S1 alone (Figure 8B). Similarly, S1-induced increased production of IL-6 was significantly (p<0.05) reduced in the presence of SKF86002 (1 μM) (Figure 8C). However, there was insignificant reduction in the production of IL-1β by SKF86002 (Figure 8D).

### S1-induced increased production of TNFα and IL-6 was reduced by TAK 242 and TLR4 siRNA

Immunofluorescence microscopy revealed that stimulation of BV-2 microglia with 100 ng/mL S1 resulted in marked increase in TLR4 expression (Figure 9A). Further experiments showed that pre-treatment with the TLR4 signalling inhibitor, TAK 242 (1 μM) resulted in significant (p<0.05) reduction in the production of TNFα and IL-6 (Figures 9B and 9C).

**Figure 8.**
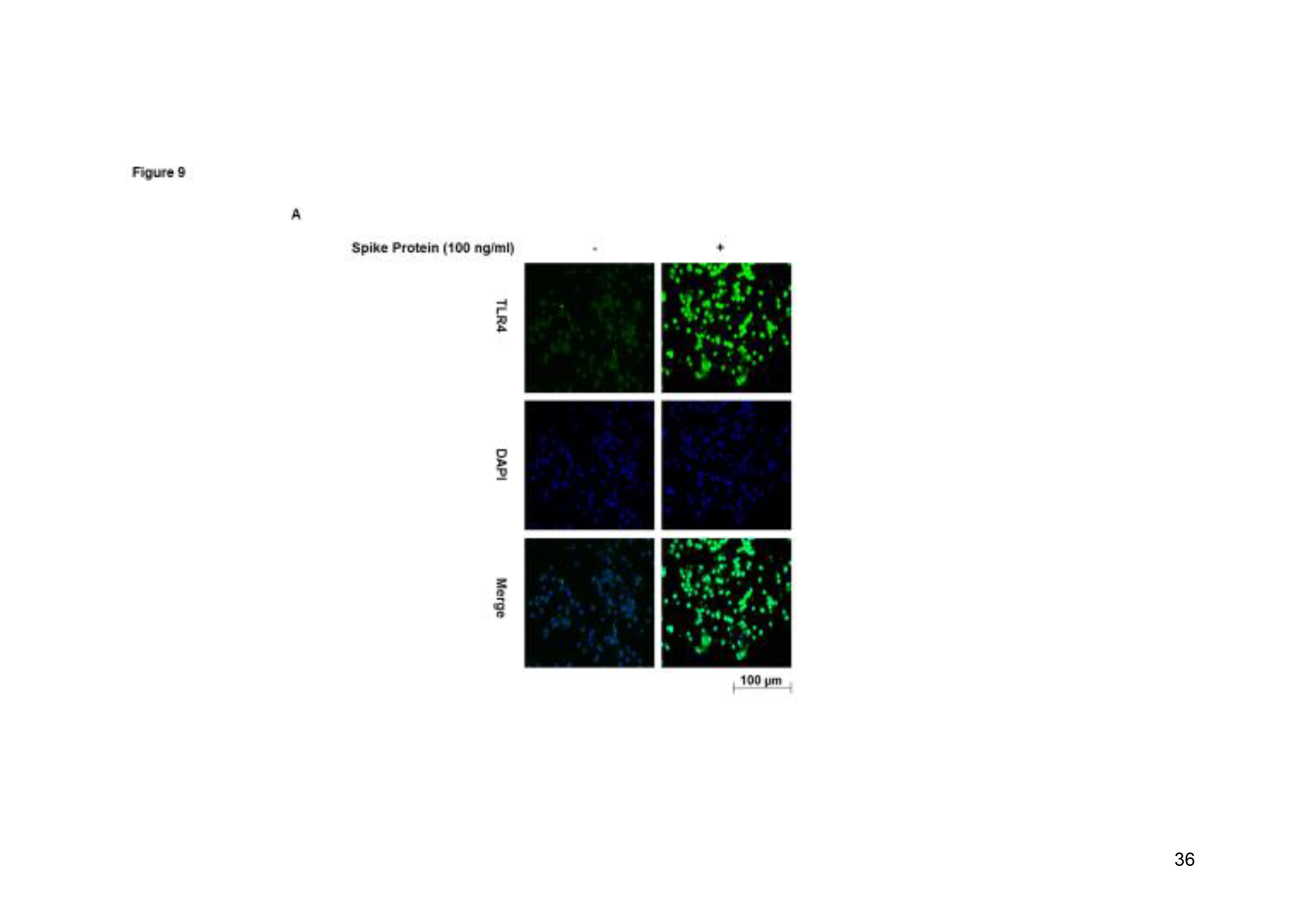
Effects of SKF 86002 (1 μM) on S1-induced increased production of TNFα (B), IL-6 and IL-1β (D) in BV-2 microglia. Culture supernatants were analysed using ELISA following stimulation for 24 h. Data were analysed using One-way ANOVA followed by a post hoc Tukey’s multiple comparison test. **p < 0.01, ***p<0.001, ****p<0.0001, ns (not significant at p<0.05), compared with untreated control, or S1 stimulation alone.

**Figure 9.**
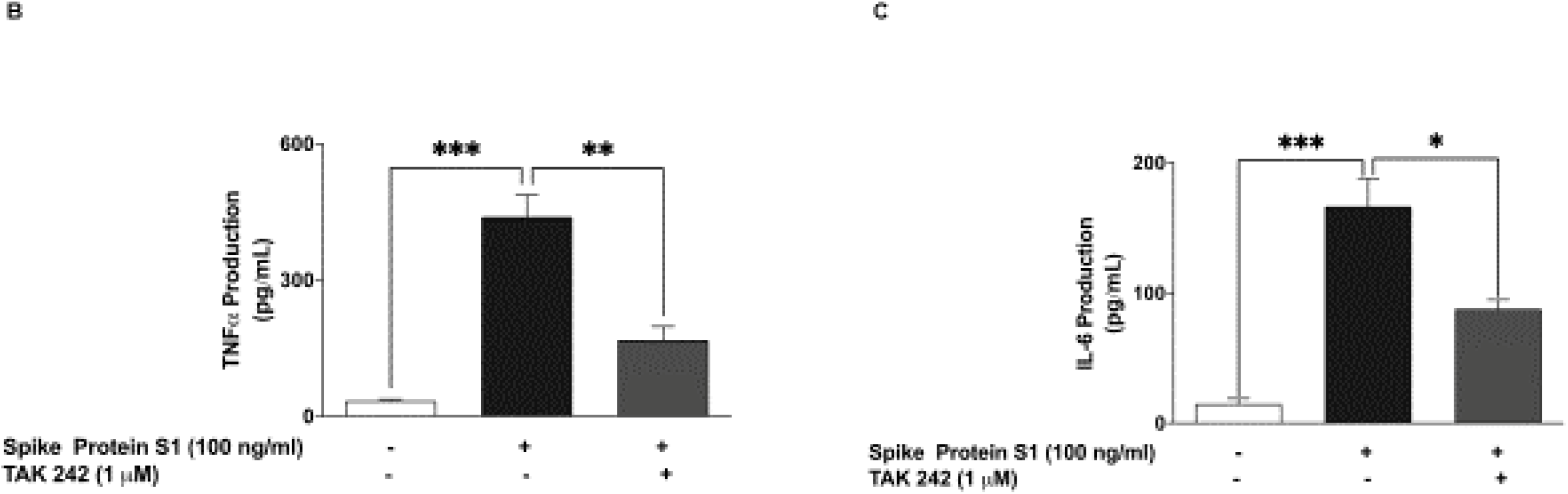
Immunofluorescence microscopy showing an increase in TLR4 protein expression following stimulation of BV-2 microglia with S1 (100 ng/mL) (A). Pre-treatment of S1-stimulated BV-2 microglia with TAK 242 (1 μM) reduced the production of TNFα (B) and IL-6 (C). Data were analysed using One-way ANOVA followed by a post hoc Tukey’s multiple comparison test. *p < 0.05, **p<0.01, ***p<0.001, compared with untreated control, or S1 stimulation alone.

**Figure 9.**
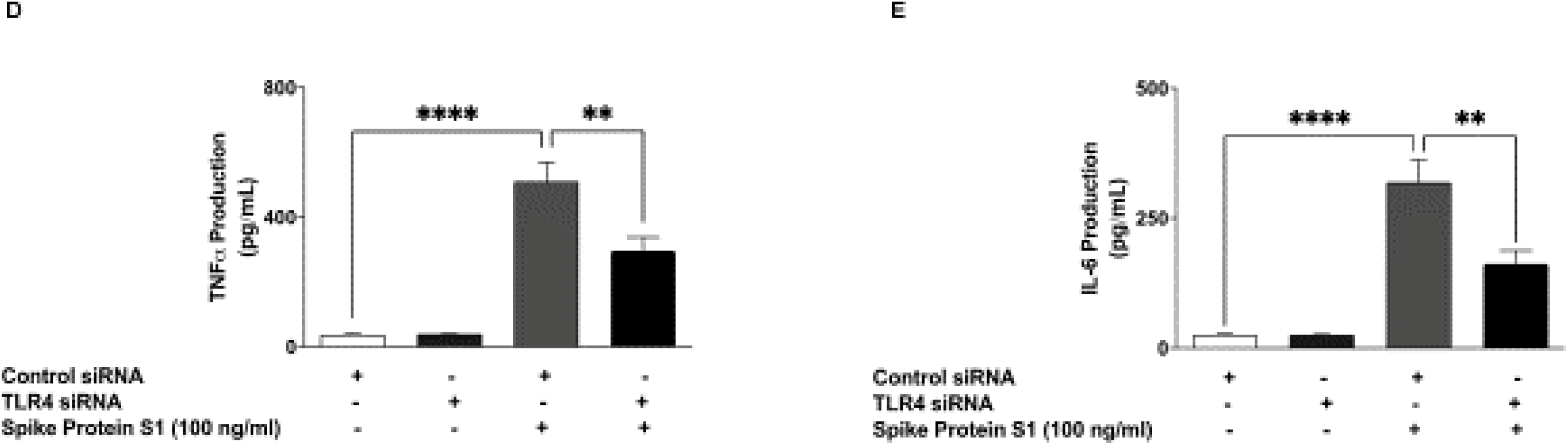
Transfection of S1-stimulated BV-2 microglia with mouse TLR4 siRNA reduced TNFα (D) and IL-6 production (E). Data were analysed using One-way ANOVA followed by a post hoc Tukey’s multiple comparison test. **p<0.01, control siRNA versus TLR4 siRNA.

Furthermore TLR4 knockdown experiments revealed that TNFα production was significantly (p<0.01) reduced in S1-stimulated BV-2 microglia transfected with TLR4 siRNA, in comparison with cells transfected with control siRNA (Figure 9D). Similarly, TLR4 knockdown resulted in reduced IL-6 production, when compared with control (Figure 9E).

## Discussion

During SARS-CoV-2 infection, viral attachment, fusion and entry into the host’s cells are facilitated by the spike proteins which protrude from the surface of mature virions, by binding to the host ACE2 protein [36, 37]. However, accumulating evidence show that in addition to facilitating its fusion to the cell membrane, the location of the spike protein on SARS-CoV-2 also makes it a direct target for host immune responses [36]. With respect to immune responses induced by SARS-CoV-2 infection, it is now established that this may involve an exaggerated inflammation, also known as cytokine storm [38–40]. Recently, we reported induction of increased release of pro-inflammatory cytokines in human peripheral blood mononuclear cells stimulated with a recombinant SARS-CoV-2 spike protein sub-unit S1, further confirming the role of the spike protein in COVID-19 cytokine storm [41].

In this study, we showed for the first time that a recombinant SARS-CoV-2 spike protein sub-unit S1 activated BV-2 microglia as demonstrated by an increase in the protein expression of Iba-1, which is mainly expressed by the microglia and increased by the activation of these cells [42, 43]. We further demonstrated that activation of BV-2 microglia by S1 resulted in increased release of TNFα, IL-6 and IL-1β, which are hallmarks of neuroinflammation. Activation of neuroinflammatory processes by the spike S1 protein was further confirmed by results showing increased iNOS-mediated production of NO by the protein in microglia. Elevated iNOS/NO has been previously linked to a wide range of CNS disorders including Alzheimer’s disease, Parkinson’s disease, multiple sclerosis, epilepsy, and migraine [44].

These are interesting outcomes, considering the roles played by microglia activation and subsequent increased release of pro-inflammatory mediators in neurological disorders. In Parkinson’s disease for example, significant increase in the expression of IL-1β was shown in the substantia nigra and frontal cortex, compared to controls [45]. Pro-inflammatory cytokine (IL-1β, TNF-α, IL-6) release has also been linked to depression-like behaviours and cognitive defects in mice [46]. Further studies are therefore needed to elucidate the link between COVID-19 mediated hyper-inflammation and the pathogenesis of neurological symptoms that have been reported to be associated with the disease.

In neuroinflammation, the NF-κB transcription factor regulates the production of multiple pro-inflammatory genes, including the pro-inflammatory cytokines such as TNFα, IL-6 and IL-1β, as well as iNOS. We further showed that activation of microglial NF-κB signalling mediates the production of pro-inflammatory mediators in the microglia by SARS-CoV-2 spike S1 protein through its ability to promote cytoplasmic phosphorylation of the p65 sub-unit and IκBα, as well as DNA binding of p65 sub-unit and NF-κB transcriptional activity. Interestingly, these effects were blocked by the potent NF-κB inhibitor, BAY11-7082. The involvement of NF-κB in S1 protein-induced neuroinflammation was further confirmed by results showing the effectiveness of BAY11-7082 in blocking S1-induced production of TNFα, IL-6 and IL-1β, and iNOS/NO in BV-2 microglia. In a similar study reported by Patra et al., SARS-CoV-2 spike protein was shown to promote IL-6 signalling through NF-κB in epithelial cells to initiate coordination of a hyper-inflammatory response [47]. To our knowledge, this is the first evidence demonstrating the role of NF-κB activation in S1-induced microglia activation.

Reports in scientific literature suggest that the NLRP3 inflammasome may be contributing to the release of cytokines such as IL-1β in SARS-CoV-2-induced hyper-inflammation [48–50]. In the microglia, activation of NLRP3 by extracellular ATP, certain bacterial toxins, crystalline and particulate matters results in caspase-1 activation which then cleaves the precursors of IL-1β and IL-18 to generate active IL-1β and IL-18, resulting in neuroinflammation and pyroptosis [51–53]. In this study, we showed that in addition to increasing IL-1β production, SARS-CoV-2 spike S1 protein activated NLRP3 inflammasome and increased caspase-1 activity in BV-2 microglia. These effects were also shown to be attenuated by CRID3 and BAY11-7082, which are known inhibitors of NLRP3. Furthermore, pre-treatment of BV-2 microglia with CRID3 prevented SARS-CoV-2 spike S1 protein-induced IL-1β-production, but not TNFα, IL-6 or NO, suggesting an involvement of the NLRP3 inflammasome activation in neuroinflammation induced by S1 in BV-2 microglia. The observed effects of BAY11-7082 on both NF-κB and NLRP3 activation in S1-stimulated BV-2 microglia reflect a potential therapeutic benefit of this compound in SARS-CoV-2 infection. Previous studies on this inhibitor of NF-κB have shown that it improved survival and reduced pro-inflammatory cytokine levels in the lungs of SARS-CoV-infected mice [54]

It has been suggested that activation of NF-κB could trigger NLRP3 through caspase-1, with subsequent release of mature IL-1β [55]. It is not clear from this study if the activation of NLRP3/caspase-1 by S1 was due to a direct effect on NLRP3 or an indirect action as a result of activating NF-κB. The role of NLRP3 inflammasome activation in S1-induced neuroinflammation therefore requires further investigation.

Our results showing activation of p38 MAPK by S1, coupled with inhibition of S1-induced increased production of TNFα and IL-6 by the p38 inhibitor (SKF 86002) suggest the involvement of the p38 MAPK signalling in its pro-inflammatory effect in BV-2 microglia. One possible explanation for this effect is the activation by S1 of upstream signals which activate both NF-κB and MAPK signalling in the microglia.

To explore the potential mechanisms involved in dual activation of NF-κB and MAPK by S1 in BV-2 microglia, we conducted experiments that showed an increase in the expression of toll-like receptor 4 (TLR4) by the protein. Interestingly, a known inhibitor of TLR4 signalling, TAK 242 inhibited S1-induced increased production of both TNFα and IL-6, which further demonstrates a potential involvement of this pattern recognition receptor (PRR) in S1-induced neuroinflammation.

Our findings seem to confirm those reported from a study by Shirato and Kizaki [56], as they demonstrated involvement of TLR4 in the induction of pro-inflammatory responses in mouse and human macrophages by SARS-CoV-2 spike protein S1 subunit. Similar observations linking TLR4 to induction of inflammation in THP-1 cells have been reported by Zhao et al. [57]. Studies to elucidate detailed modulatory effects of SARS-CoV-2 spike protein S1 subunit on microglia TLR4 signalling are required in order to provide better understanding of the impact of SARS-CoV-2 infection on the brain.

The obtained from these experiments have provided the first evidence demonstrating activation of BV-2 microglia by SARS-CoV-2 spike S1 protein, resulting in the production pro-inflammatory mediators TNFα, IL-6, IL-1β and NO. We propose that induction of neuroinflammation by this protein in the microglia is mediated through activation of NF-κB and p38 MAPK, possibly as a result of TLR4 activation. These results contribute to our understanding of some of the mechanisms involved in CNS pathologies of SARS-CoV-2.

## Author contributions

**Olumayokun A Olajide**: Conceptualisation, Methodology, Investigation, Writing - Original Draft, Writing- Review & Editing, Project administration. **Victoria U Iwuanyanwu**: Investigation. **Oyinkansola D Adegbola**: Investigation. **Alaa A Al-Hindawi**: Investigation.

## Compliance with ethical standards

### Research involving Human Participants and/or Animals

Not applicable.

### Informed consent

Not applicable.

### Consent to participate

Not applicable.

### Consent for Publication

Not applicable.

## Acknowledgments

Not applicable.

## Data availability statement

The datasets generated during and/or analysed during the current study are available from the corresponding author on reasonable request.

