## Supplementary Data for "SARS-CoV-2 spike glycoprotein S1 induces neuroinflammation in BV-2 microglia"

Effects of 10, 50, 100, 500 and 1000 ng/mL of S1 on TNFα production following stimulation of BV-2 microglia for 1, 3, 6, 12 and 24 h

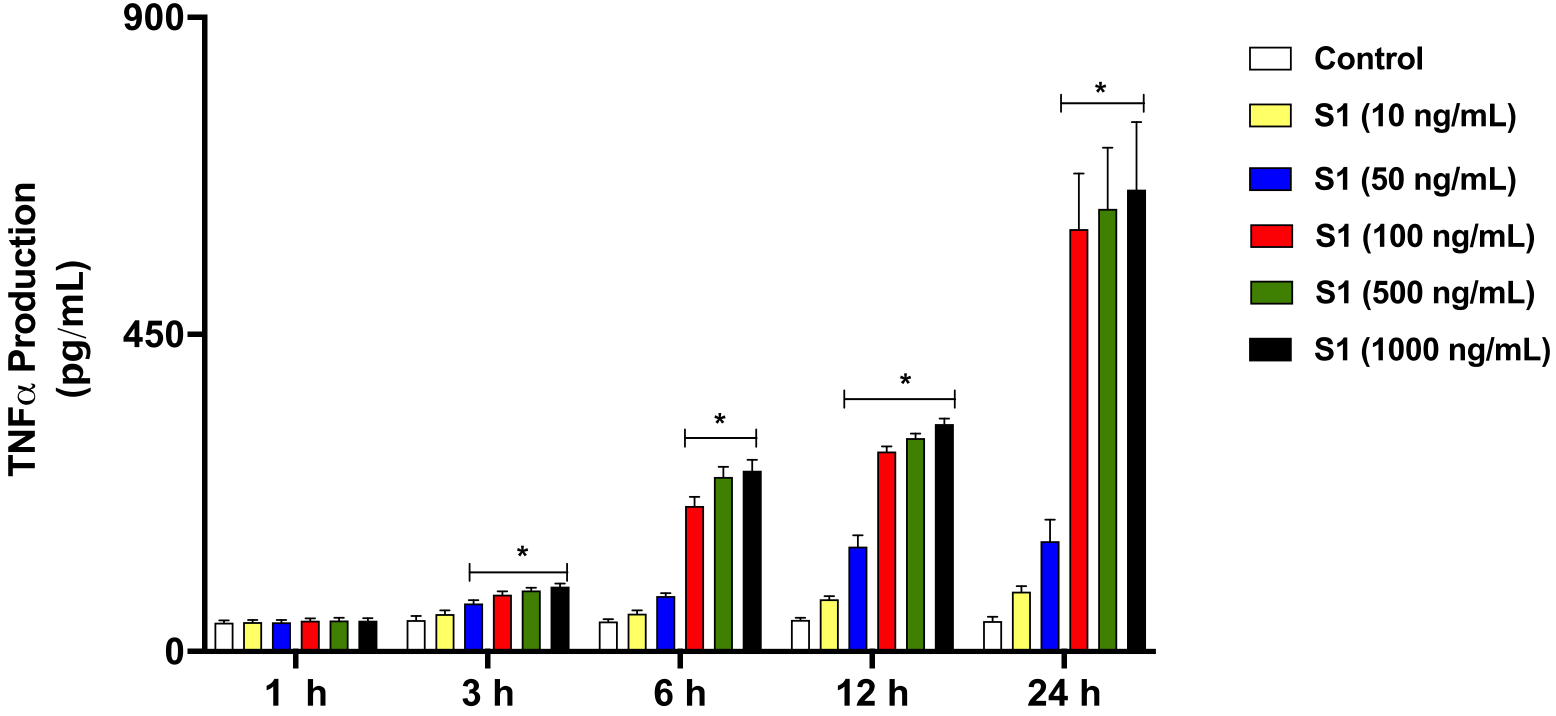
